# Beyond the seed: structural basis for supplementary microRNA targeting

**DOI:** 10.1101/476960

**Authors:** Jessica Sheu-Gruttadauria, Yao Xiao, Luca F. R. Gebert, Ian J. MacRae

## Abstract

microRNAs (miRNA) guide Argonaute proteins to mRNAs targeted for repression. Target recognition occurs primarily through the miRNA seed region, composed of guide (g) nucleotides g2–g8. However, nucleotides beyond the seed are also important for some known miRNA-target interactions. Here, we report the structure of human Argonaute2 (Ago2) engaged with a target RNA recognized through both miRNA seed and supplementary (g13–g16) regions. Ago2 creates a “supplementary chamber” that accommodates up to 5 miRNA-target base pairs. Seed and supplementary chambers are adjacent to each other, and can be bridged by an unstructured target loop of 1–15 nucleotides. Opening of the supplementary chamber may be constrained by tension in the miRNA 3' tail as increases in miRNA length stabilize supplementary interactions. Contrary to previous reports, we demonstrate optimal supplementary interactions can increase target affinity >20-fold. These results provide a mechanism for extended miRNA-targeting, suggest a function for 3' isomiRs in tuning miRNA targeting specificity, and indicate that supplementary interactions may contribute more to target recognition than is widely appreciated.

## Introduction

MicroRNAs (miRNA) are small non-coding RNAs that associate with Argonaute proteins and pair with messenger RNAs (mRNA) targeted for repression. In miRNA targeting, the seed region (nt g2-g7 or g2-g8, from the miRNA 5′ end) is the primary determinant for targeting efficacy and specificity, with over 80% of miRNA-target interactions occurring through seed-pairing (Grosswendt, Filipchyk et al., 2014). This observation is unsurprising, as the seed region is the most evolutionarily conserved portion of the miRNA (Brennecke, Stark et al., 2005, Krek, Grun et al., 2005, Lewis, Burge et al., 2005, Lewis, Shih et al., 2003), and indeed, seed-pairing alone is sufficient to induce significant levels of target repression (Brennecke et al., 2005, Doench & Sharp, 2004, Lim, Glasner et al., 2003). However, it has been recurrently demonstrated that additional regions of the miRNA outside of the seed can contribute to target recognition (Broughton, Lovci et al., 2016b, Grimson, Farh et al., 2007a, Moore, Scheel et al., 2015, Salomon, Jolly et al., 2015, Wee, Flores-Jasso et al., 2012).

Beyond the seed, nucleotides in the supplementary region (g13-g16) have been implicated in miRNA targeting in numerous studies. It is thought that supplementary nucleotides can enhance or reinforce recognition of seed matched targets (Brennecke et al., 2005, Grimson et al., 2007a), although conservation of such sites appears to be relatively rare (Friedman, Farh et al., 2009). In accordance with this idea, biochemical studies indicate that supplementary pairing has only a modest influence (~2 fold) on target affinity for certain miRNA sequences (Salomon et al., 2015, Wee et al., 2012). Nevertheless, it has been suggested that 3′ pairing may compensate for weak or mismatched seed-pairing, or may even function in the absence of seed complementarity (Brennecke et al., 2005, Grimson et al., 2007a). These so-called 3′ compensatory sites have been widely identified in several high-throughput crosslinking immunoprecipitation studies (Chi, Hannon et al., 2012, Grosswendt et al., 2014, Hafner, Landthaler et al., 2010, Helwak, Kudla et al., Loeb, Khan et al., 2012), although recent computational re-analysis indicates that while these sites may interact with miRNAs, they do not correlate with induced repression (Agarwal, Bell et al., 2015). It remains to be determined, however, whether these sites serve other functional purposes or contribute to silencing under certain biological contexts.

Most mature miRNAs are broadly categorized into families that share identical seed sequences but often contain divergent 3′ regions (Bartel, 2018). Many models suggest that miRNAs within families are largely redundant and likely associate with the same target RNAs, depending on their relative expression levels. In contrast with this idea, human cardiomyocytes express both miRNA family members miR-25 and miR-92, but only miR-25 is capable of regulating SERCA2a transcripts in vivo (Wahlquist, Jeong et al., 2014). It has also been shown that down-regulation of certain miRNAs leads to a significant phenotypic response, despite the presence of other family members (Bartel, 2018). Recent work has directly demonstrated that pairing to the 3′ end of the miRNA can play a substantial role in targeting specificity amongst miRNA family members (Broughton et al., 2016b, Moore et al., 2015). Together, these findings highlight the ambiguity surrounding the redundancy within miRNA families.

Lastly, it has also been shown that Hepatitis C viral RNA is bound by Ago2-miR-122, which is thought to be involved in genome stabilization and infectivity (Shimakami, Yamane et al., 2012). This process requires supplemental base pairing in addition to the seed (Machlin, Sarnow et al., 2011). These findings further indicate that seed sequence alone is not always sufficient for determining functional target interactions.

miRNA target prediction algorithms still face substantial challenges, despite remarkable advances in technology and validation (Agarwal et al., 2015, Hausser & Zavolan, 2014). Current approaches rely heavily on identification of seed-matched sites within annotated 3′ UTRs. However, because the seed region is only composed of 6 to 7 contiguous nucleotides, such sites appear often in the transcriptome and are not always functionally responsive to the associated miRNA (Baek, Villen et al., 2008, Selbach, Schwanhausser et al., 2008, Yue, Liu et al., 2009). Therefore, seed-matched target identification is often combined with additional predictive strategies, including evolutionary conservation and positional context within the 3′ UTR (Agarwal et al., 2015, Grimson et al., 2007a). However, even with these major advancements, false positives and negatives are still prevalent, indicating potential for further improvements in predictive algorithms.

A current limitation in understanding the miRNA 3′ region arises from a lack of structural and biochemical information exploring to how Argonaute accommodates extended pairing. To date, all structural studies of human Argonaute have been limited to targets that interact with the extended seed region (Schirle, Sheu-Gruttadauria et al., 2015, Schirle, Sheu-Gruttadauria et al., 2014). Here, we present the crystal structure of human Argonaute2 (Ago2) bound to a miRNA engaged with a target through seed and supplementary base pairing. The structure shows supplementary pairing occurs in a defined supplementary chamber that can accommodate up to 5 miRNA-target base pairs within the central cleft of Ago2. The supplementary chamber is physically separated from the chamber that houses the seed-paired duplex (seed chamber) by a central gate in Ago2, which prevents targets from interacting with the miRNA central region (g9–g11). Due to compaction of the miRNA central region within Ago2, seed and supplementary chambers are in close proximity to each other and can be bridged by as few as 1–2 target nucleotides. Opening of the supplementary chamber to allow miRNA-target pairing requires movements of the Ago2 PAZ domain, which appears to be tethered by interactions with the miRNA 3′ end. Consistent with this model, extending miRNA length substantially increases the stability and versatility of supplementary interactions. Moreover, contrary to previous reports, biochemical measurements show that supplementary interactions can enhance the affinity of seed-paired targets by more than an order of magnitude, and for strong (GC-rich) supplementary interactions seed and supplementary can be bridged by target loops up to 15 nt in length. The combined results reveal the structural basis for supplementary miRNA-target interactions, provide structure-guided criteria for better defining and predicting supplementary interactions, and suggest an unexpected role in differential targeting by miRNAs of varying 3′ lengths, called isomiRs.

## Results

### An unanticipated crystal structure indicates Ago2 avoids central pairing

Although this study ultimately revealed the structural basis for supplementary miRNA target interactions, our crystallization efforts were originally aimed at determining the structural basis for target RNA cleavage, or “slicing”, by small RNAs. A catalytically inactive mutant Ago2 (D669A) was used to avoid target cleavage. The sequence of the guide RNA used corresponded to a 21 nt isoform of miR-122. Target RNA design was based on the observation that pairing to the guide tail region (g17–g21) is not necessary for efficient cleavage (Ameres, Martinez et al., 2007, Chang, Yoo et al., 2009, De, Young et al., 2013, Haley & Zamore, 2004, Wee et al., 2012), and that extensive pairing, especially in the 3′ tail, induces a phenomenon called “unloading,” in which the guide:target duplex is released from purified samples of Ago2 (De et al., 2013, Jo, Shin et al., 2015, Park, Shin et al., 2017). We therefore initially screened for a slicing-competent conformation using a target RNA that was fully complementary to positions g2-g16 of miR-122. A t1-adenosine was included to increase target affinity (Schirle et al., 2015).

Surprisingly, however, the complex crystallized in a conformation that avoids central pairing entirely, and instead appeared to be more representative of a complex with target pairing to the seed and supplementary regions of the miRNA (Fig. 1B). This observation indicates that, even when central pairing is available, Ago2 molecules can maintain a stable conformation in which central pairing is avoided. A caveat to this conclusion is that the structure contains Ago2 with a point mutation (D669A) that abolishes slicing activity, which otherwise would likely have led to target cleavage over the incubation period (7-14 days) required for crystallization. However, the observed structure is consistent with the previous observation that Ago2 often binds perfect complement target RNAs without cleaving them (Wee et al., 2012).

**Fig. 1.**
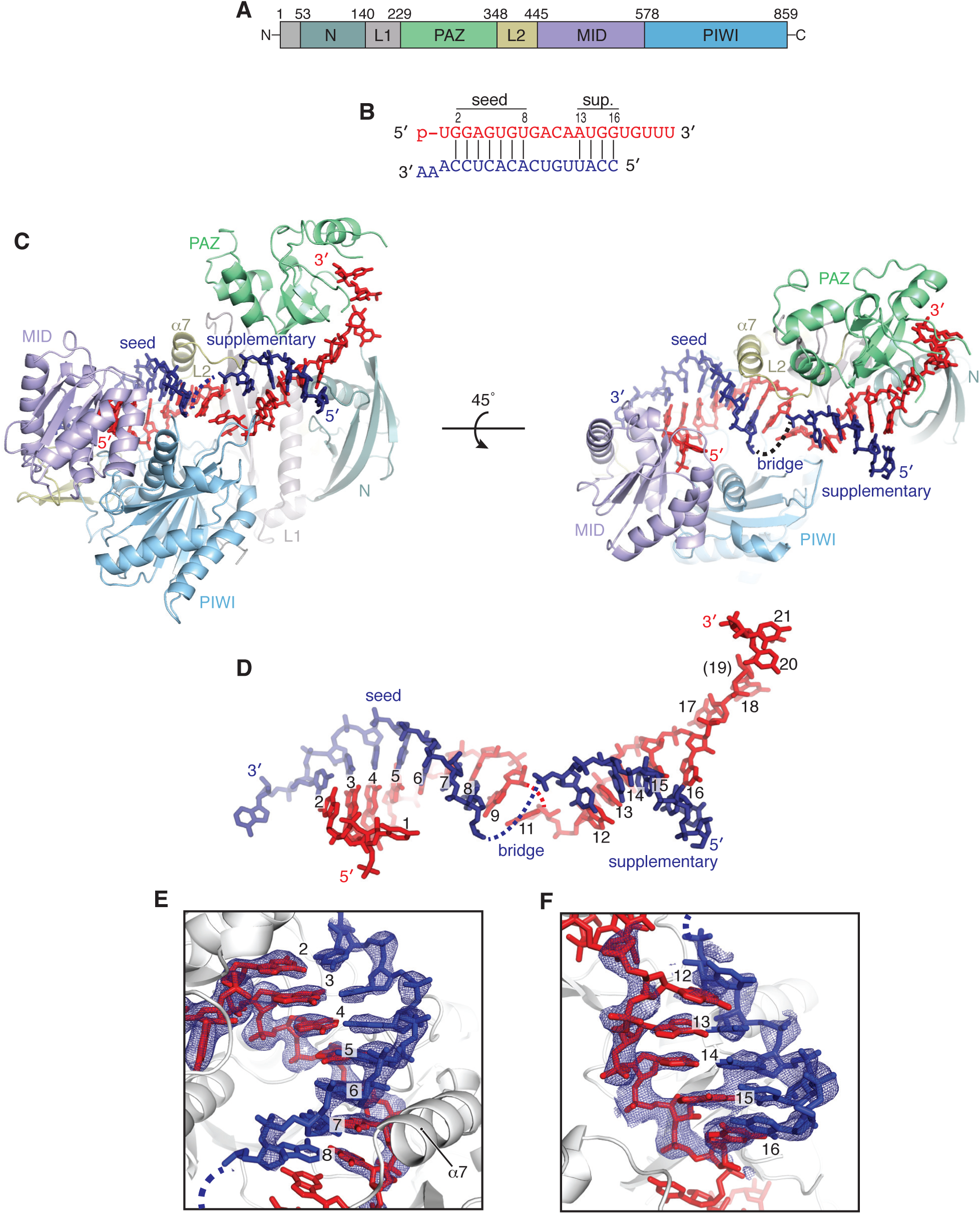
Overview of seed plus supplementary structure. **A.** Schematic of Ago2 primary sequence with domains colored as in figures throughout manuscript. **B.** Schematic of crystalized miRNA and target sequences. Vertical lines indicate base pairs observed in structure. miRNA colored red; target RNA colored dark blue (here and throughout manuscript). **C.** Front and top views of Ago2 bound to seed plus supplementary-paired target RNA. **D.** miRNA-target duplex with miRNA nucleotides numbered from 5′ end (Ago2 removed for clarity). Dashed line indicates disordered nucleotides in bridge region. **E.** Close-up of seed-paired region of miRNA-target duplex. Fo-Fc electron density map calculated after omitting nucleotides around the seed region (miRNA and target at positions 1–9), contoured at 3.0 sigma, shown as blue mesh. **F.** Close-up of supplementary-paired region of miRNA-target duplex. Fo-Fc electron density map calculated after omitting nucleotides in the supplementary region (miRNA and target at positions 12-16), contoured at 2.0 sigma, shown as blue mesh.

### Structure of human Ago2 in complex with a seed plus supplementary-paired target RNA

An overview of the seed plus supplementary-paired structure reveals the miRNA and target RNA form two discontinuous duplexes: one composed of the seed region, and a secondary duplex in the supplementary region (Fig. 1C-D). Within the seed region, the target RNA pairs to nucleotides g2-g8, with the t1-adenosine sequestered within the t1-binding pocket, as reported in seed-paired structures. miRNA:target pairing ends after nucleotide g8. Electron density for t8 (the target nucleotide opposite g8) is weaker than the rest of the seed-paired target nucleotides, indicating a higher degree of mobility (Fig. 1E). After the seed, the miRNA deforms away from A-form, with the g9 nucleobase adopting the syn-conformation and stacking against both nucleobases in the g8-t8 base pair to cap the seed-paired duplex (Fig. 1E). Nucleotide g10 is disordered, and density for the base of g11 was difficult to interpret. g11 immediately precedes the narrowest point in central cleft, after which g12 emerges within the supplemental chamber. Due to the narrowing of the central cleft and the position of loops in the PIWI (residues 602-608) and L2 (residues 351-358) domains (henceforth called the PIWI Loop and L2 Loop, respectively), target pairing is prevented to all central nucleotides and instead re-initiates at position g13 (Fig. 1C). The crystallized conformation, therefore, prevents topological winding of the miRNA:target duplex within the central cleft.

Target nucleotides t9–t11 (the nucleotides opposite miRNA g9–g11) form a bridging region between seed and supplementary duplexes and are disordered in the observed structure. Steric clashes between the L2 Loop and the backbone of t12 prevent formation of a base pair with g12, which is exposed and well positioned for target pairing (Fig. 1D). t12 is instead shifted to partially stack beneath g12, thereby capping the supplementary duplex (Fig 1F). In the observed conformation, the supplementary chamber is only wide enough to allow the target RNA to pair continuously to positions g13-g16. Notably, a minor shift in the position of the L2 Loop would allow up to five contiguous supplementary base pairs. The miRNA 3′ tail (g17–g21) is threaded through a narrow gap between the N and PAZ domains (the N-PAZ channel), and the miRNA 3′ end is bound in the PAZ 3′ binding pocket (Ma, Ye et al., 2004) (Fig 1C). The 5′ end of t16 is outside of the supplementary chamber and completely solvent exposed, indicating target nucleotides 5′ of the supplementary region do not impact interactions with the miRNA (Fig. 1C).

### A structural model for supplementary targeting

Previously, we reported structures of human Ago2 bound to a seed-paired target RNA (Schirle et al., 2014). These structures provided a model for miRNA targeting, wherein seed pairing results in a conformational shift in the protein that widens the N-PAZ channel to reveal the supplementary chamber. This movement is coupled with rearrangement of the miRNA such that the nucleobases of g12-g16 stack against each other, resembling the seed-region prior to target association. However, the Watson-Crick faces of nucleotides g12–g14 are eclipsed by the L2 Loop and PIWI Loop in seed-only paired structures. Therefore, to accommodate contiguous pairing to the supplementary region, we predicted a more substantial conformational change would be necessary than observed with seed pairing alone (Schirle et al., 2014).

Side-by-side comparison of the seed-paired and seed plus supplementary-paired Ago2 structures reveals the conformational shifts necessary for pairing at positions g13-g16 (Fig. 2A). Moving from the seed-only conformation, the L2 Loop and PIWI Loop shift away from each other by ~7Å, and thereby provide space necessary for the target RNA to pair with the miRNA at positions g13 and g14 (Fig. 2A). This movement allows repositioning of the 3′ half of the miRNA, which adopts a more extended conformation due to movement of the PAZ domain (connected to the L2 loop and helix-7), and rises within central cleft to fully expose nucleotides g12-g16 for pairing (Fig. 2B).

**Fig. 2.**
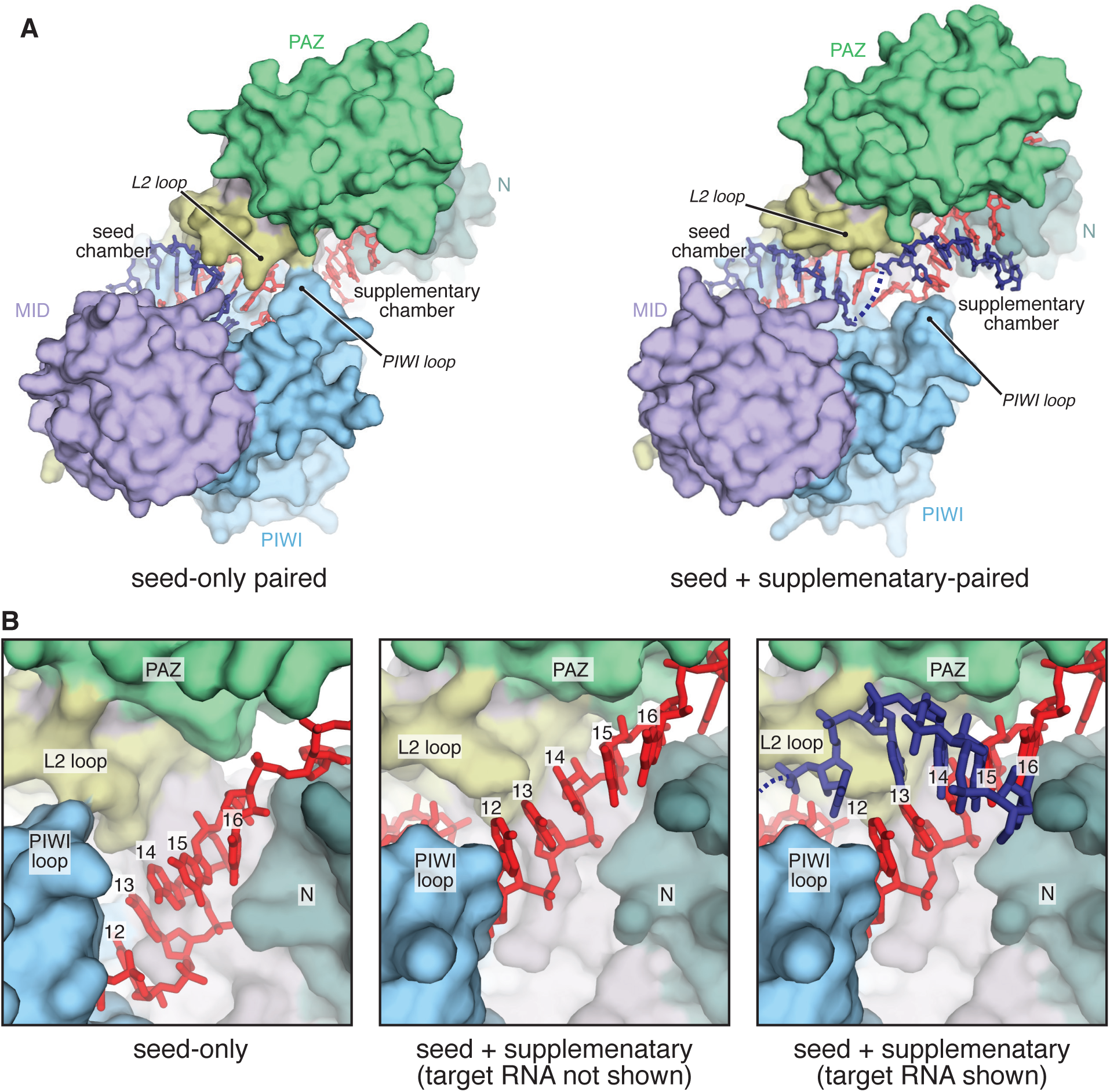
Side-by-side comparison of seed-only and seed plus supplementary-paired Ago2 structures. **A.** Surface representation of seed-only paired (left, PDB 4Z4D) and seed plus supplementary-paired (right) Ago2 structures illustrates opening of supplementary chamber. RNAs shown as sticks. Seed and supplementary chambers indicated. **B.** Close-up view of supplementary chamber in seed-only (left) and seed plus supplementary-paired (middle and right) structures illustrates movement of miRNA upon supplementary interactions.

Superimposing the seed-only and seed plus supplementary-paired structures reveals conformational differences associated with supplementary pairing. Differences within the Ago2 MID-PIWI lobe are restricted to the PIWI Loop, which shifts ~3 Å between seed-only and seed plus supplementary structures (Fig. 3A). Likewise, the N-PAZ lobes are similar in both structures, though N and PAZ domains are slightly twisted in opposite directions about the central L1 domain (Fig 3B). Opening of the central cleft to accommodate supplementary pairing is primarily achieved though the action of a hinge composed of β-strand-1 (residues R36–A42) near the N-terminus, and β-strand-19 (residues T406–V412) of the L2 domain, which form the outer two strands of the central β-sheet in the PIWI domain, and resides at the connection point between the N-PAZ and MID-PIWI lobes (Fig. 3C).

**Fig. 3.**
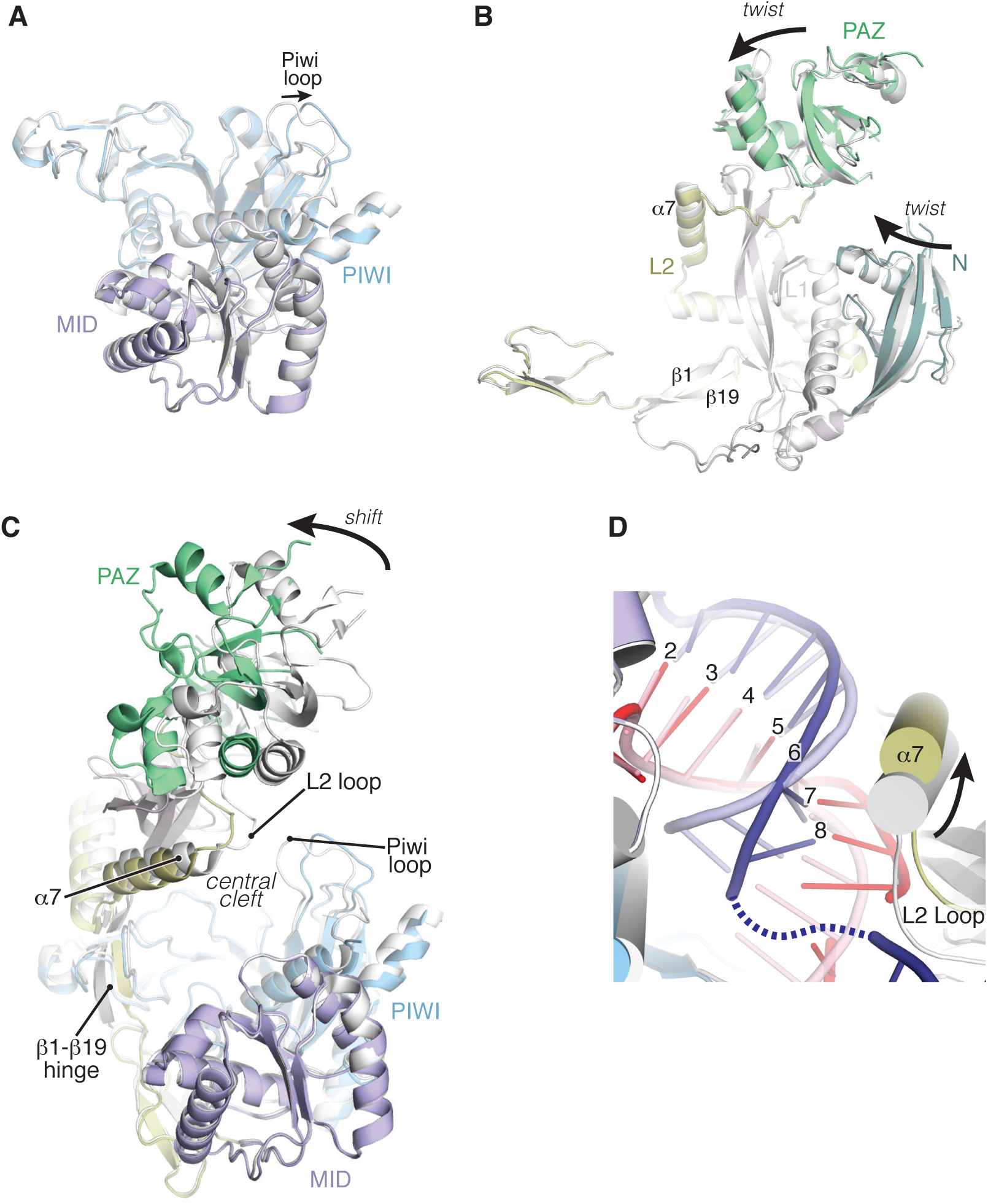
Structural analysis of supplementary pairing. **A.** Superposition of MID-PIWI lobes from seed-only (gray) and seed plus supplementary-paired (colored) structures shows only the PIWI Loop changes substantially between upon supplementary interactions (RNAs not shown for clarity). **B.** Superposition of N-PAZ lobe reveals minor twists in N and PAZ domains relative to L1, but overall structures are very similar between seed-only and seed plus supplementary-paired structures. **C.** Superposition of entire Ago2 structures reveals major differences are restricted to rigid body movement of the N-PAZ lobe relative to the MID-PIWI lobe. Movement centers around the β1-β19 hinge. **D.** Superposition of seed chambers in the seed-only paired (gray/semi-transparent) and seed plus supplementary-paired (colored) structures. Helix-7 (α7) contacts miRNA and target in both structures despite conformational shifts to accommodate opening of the supplementary chamber.

The position of the seed-paired miRNA-target duplex also shifts upon addition of supplementary interactions. Across both structures, nucleotides g2-g6 remain largely stationary, with the PIWI domain interrogating the minor groove of the miRNA-target duplex in both complexes. However, after nucleotide g6, the seed-paired duplex shifts ~3Å away from the MID-PIWI lobe of the protein. The shift is accommodated by an equal movement in helix-7 of Ago2, which remains docked within the minor groove of the seed-paired duplex (Fig. 3D). Helix-7 is directly connected to the L2 Loop, which both move as a rigid body with the PAZ domain ~3Å away from the central cleft. Combined with the positional shift in the PIWI Loop described above, these conformational changes allow Ago2 to remain engaged with the seed-paired target while providing space necessary for miRNA-target interactions in the supplementary chamber.

### Seed and supplementary duplexes can be bridged by a target loop of variable size

Based on a global analysis of miRNA target sites, the most efficacious position for supplementary-paired region on a target RNA is directly opposite the guide RNA, although offsets of ~2 nt appear to be tolerated (Grimson et al., 2007a). These findings predict a loop of ~1-5 nt bridging between the seed and supplementary duplexes (Grimson et al., 2007a). In the crystallized structure, compaction of the miRNA central region brings the seed and supplementary duplexes within close proximity of each other (Fig. 1D). This proximity may explain why relatively short loops (1-2 nt) in targets can functionally bridge the seed and supplementary regions. Notably, although the bridging nucleotides were disordered in our structures, and thus could not be modeled, the positions of the seed and supplementary duplexes indicate the bridging loop must be contained within the relatively narrow gap between the N-PAZ and MID-PIWI lobes of Ago2 (Fig. 4A). We hypothesized that this physical constraint may decrease the entropic cost in bridging seed and supplementary duplexes and thereby allow a larger variety of sizes of functional bridging loops than is currently appreciated. The contribution of bridging loop length to target affinity, however, has not been interrogated.

**Fig. 4.**
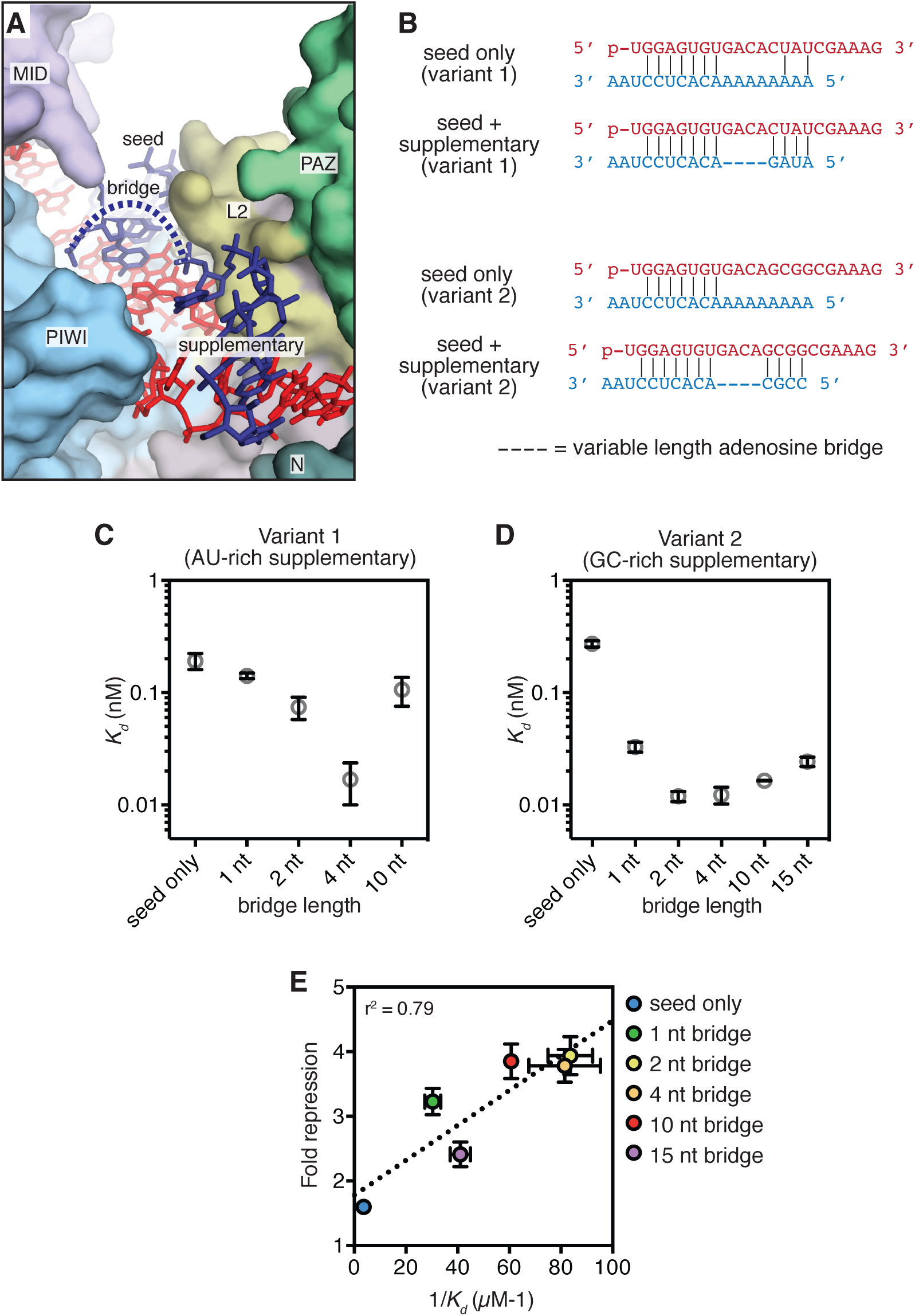
Impact of seed-supplementary bridging region on target affinity. **A.** Overview of central cleft shows path of target RNA. Seed, bridge, and supplemental regions indicated. **B.** Schematic of pairing between miRNAs (loaded into Ago2) and targets used in equilibrium binding experiments. **C.** Dissociation constants (*K*_*d*_) of Ago2-miRNA complexes (containing an AU-rich supplementary region) binding to target RNAs with bridging regions of various lengths. **D.** Dissociation constants (*K*_*d*_) of Ago2-miRNA complexes (containing an GC-rich supplementary region) binding to target RNAs with bridging regions of various lengths. **E.** Fold repression of a luciferase reporter (relative to no target site control) plotted versus affinity (1/*K*_*d*_) values measured for the same sites in vitro. All plotted data are the means of at least three independent replicates. Error bars indicate SEM.

We measured the affinity of Ago2-miRNA for target RNAs that contained seed and supplementary-paired regions separated by various numbers of bridging nucleotides (Fig. 4 and EV1). Care was taken to design target sequences that do not contain internal structure and have complementarity restricted to the miRNA seed and supplementary regions (Fig. 4B). We first examined miRNA-target combinations with a relatively weak (AU-rich) supplementary sequence at positions g13–g16. As expected, targets capable of supplementary pairing had higher affinities for Ago2-miRNA than the seed-only matched control (Fig. 4C). Target affinity varied with bridging loop length and a 4 nt bridging loop provided the highest affinity of all targets tested (>10-fold greater than the seed-only target). Notably, even a target with an unstructured bridging loop of 10 nt displayed a modest (1.9-fold), but significant (p=0.028, two-tailed, unpaired Student’s t-test) increased affinity compared to the seed-only control.

We also examined the effects of bridging loop length using a stronger (GC-rich) supplementary sequence (Fig. 4D). Again, a bridging region closely matching the miRNA (i.e. 2–4 nt) provided the greatest increase in target affinity (>20-fold over the corresponding seed-only target). However, bridges ranging in size from 1 to 15 nt also showed marked (i.e. >10-fold) increases in affinity. Moreover, in an optimal case (4 nt bridge, complementarity at g13–g14) even two GC supplementary base pairs were sufficient to increase target affinity >3.5-fold (p<0.0001, two-tailed, unpaired Student’s t-test) (Fig. EV2).

The finding that a 4 nt bridge provides optimal target affinity is consistent with global analyses of 3′ supplementary target site efficacy and conservation (Grimson, Farh et al., 2007b). This connection suggests that increased target affinity associated with supplementary interactions may translate into enhanced repression in mammalian cells. To test this hypothesis, we measured repression of a luciferase reporter bearing a single target site to variants of miR-122 transfected into HEK 293 cells (Fig. EV3). For an optimal target (GC-rich supplementary sequence, 4 nt bridging loop) we found supplementary interactions increased repression 2-fold over the seed-only target reporter. Additionally, as observed in binding assays, an unstructured bridging loop of up to 15 nt supported an observable increase in repression (Fig. 4E). Plotting in vitro target affinity (1/*K*_*d*_) as a function of luciferase repression of revealed a modest correlation (r^2^ = 0.79).

### Stabilized supplementary interactions by 3′ isomiRs

The large increase (>20-fold) in target affinity afforded by optimal supplementary pairing reported here differs from more modest effects reported previously: ~2-fold for let-7 (Wee et al., 2012), and ~7-fold for miR-21 (Salomon et al., 2015). Part of this discrepancy may be attributed to differences in the miRNA seed and supplementary sequences (Salomon et al., 2015). Additionally, we noticed that previous reports used 21 nt miRNAs, while the binding data described above were conducted with 22mers.

Inspection of the seed plus supplementary-paired crystal structure, which contains a 21 nt miRNA, reveals that the miRNA must adopt an extended conformation to be available for pairing in the supplementary region while still maintaining interactions with the miRNA 3′ binding pocket in the PAZ domain (Fig. 5). This extended conformation appears to be imposed by the position of the PAZ domain and L2 loop, which must shift away from the central cleft to allow space for the supplementary-paired target RNA (Fig 3C). Thus, movements of the PAZ domain are necessary for opening the supplementary chamber to enable supplementary interactions, but may also inhibit supplementary interactions by creating tension in the miRNA 3′ tail. Indeed, in the crystallized conformation the L2 loop prevents target pairing at g12 (Fig. 2B), which would be possible if the supplementary chamber were to open only slightly more. A clear prediction of this model is that relieving tension in the miRNA 3′ tail, by increasing miRNA length, should stabilize supplementary interactions with target RNAs.

**Fig. 5.**
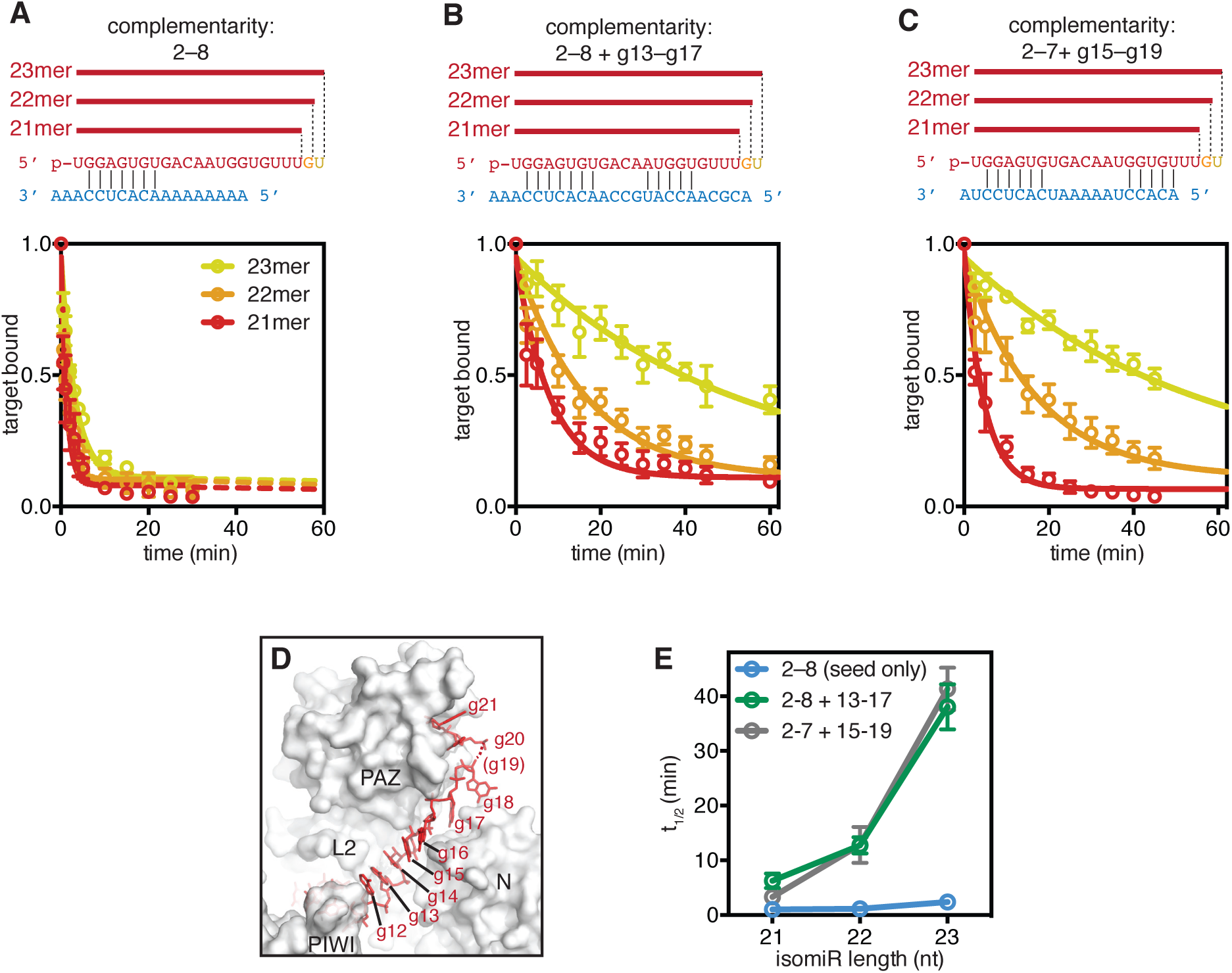
Stabilization of supplementary interactions with increasing 3′ miRNA length. **A.** Top: schematic of miR-122 isomiRs paired with a target complementary to the seed region only. Bottom: release of the seed-paired target from Ago2-miR complexes was assessed by monitoring the fraction of the total target RNA bound as a function of time. **B.** Top: schematic of miR-122 isomiRs paired with a target complementary to the seed and supplementary (g13–g17) regions. Bottom: fraction target bound to the Ago2-miR complexes as a function of time. **C.** Top: schematic of miR-122 isomiRs paired with a target complementary to the seed and shifted supplementary (g15–g19) regions. Bottom: fraction target bound to the Ago2-miR complexes as a function of time. **D.** Structure of the miRNA 3′ tail suggests tension in the miRNA may restrict PAZ domain movments and opening of the supplementary chamber (target RNA not shown for clarity). **E.** Release half times (t_1/2_) of target RNAs in panels A-C plotted against isomiR length. All plotted data are the means of at least three independent replicates. Error bars indicate SEM.

We tested this prediction by measuring the release rates of target RNAs from Ago2 loaded with 21mer, 22mer, or 23mer isomiRs of miR-122. The target used had complementarity to the seed (g2–g8) and extended supplementary (g13–g17) regions. In accordance with our model, we found that increasing miRNA length correlated with stability of the Ago2-miRNA-target ternary complex (Fig. 5B). Ago2 loaded with a 21mer guide released the target with a half-time of 5.5 minutes while Ago2 loaded with a 22mer or 23mer retained the target 2-fold or 6.5-fold longer (t_1/2_ = 11 and 37 minutes for 22mer and 23mer, respectively). isomiR effects were less pronounced for a seed-only paired target, which was released much faster in all cases (t_1/2_ = 0.9, 1.0, and 2.1 minutes for 21mer, 22mer, and 23mer, respectively) (Fig. 5A). Moreover, a related target with 3′ shifted supplementary complementarity (to g15–g19) displayed marked isomiR effects, with stability of the ternary complex increasing >10-fold with increasing length (t_1/2_ = 3.2, 12, and 38 minutes for 21mer, 22mer, and 23mer, respectively) (Fig. 5C). These observations demonstrate that 3′ tail length dictates the stability of supplementary interactions, and raise the possibility that additions to the miRNA 3′ end may allow Ago2 to expand the nucleotides available for supplementary target interactions.

### The supplementary duplex is mobile

The effects of 3′ tail length suggest that the supplementary duplex may be mobile within the supplementary chamber. Indeed, in contrast to the seed-region, the protein makes only a handful of contacts to the supplementary duplex. A603 and G604 (in the PIWI Loop) make hydrophobic/van der Waals interactions with the nucleobase of g12, and Ile-353 (in the L2 Loop) inserts into the minor grove of the 5′ end of the supplementary duplex (with respect to the miRNA) (Fig. 6A). Salt-linkages between the miRNA phosphate backbone and R68 and R635 are also observed (Fig. 6A). Notably, the supplementary chamber contains several additional arginine residues (R97, R167, R179 and R351) that are near the supplementary duplex but not close enough to directly contact in the crystallized structure (Fig 6A), indicating additional conformations may exist in solution. We therefore suggest that the supplementary duplex may be relatively mobile within the supplementary chamber and may even experience a degree of rotation around its helical axis while remaining bound to Ago2. Consistent with this notion, temperature factors of atoms in the supplemental duplex are substantially higher than atoms in the immobilized seed-paired duplex (Fig. 6B). In the case of longer 3′ isomiRs, we suggest that this mobility may allow miRNA nucleotides 3′ of g12–g13 to slide into the supplementary chamber and make productive interactions with target RNAs. Flexibility would also allow the supplementary duplex to interrogate for further complementarity either towards the central region or the 3′ end of the guide RNA. Taken together with the notion that pairing can occur in this region without topologically costly conformational changes, we suggest that supplementary interactions may serve as an initial nucleation site for 3′ and/or central pairing.

**Fig. 6.**
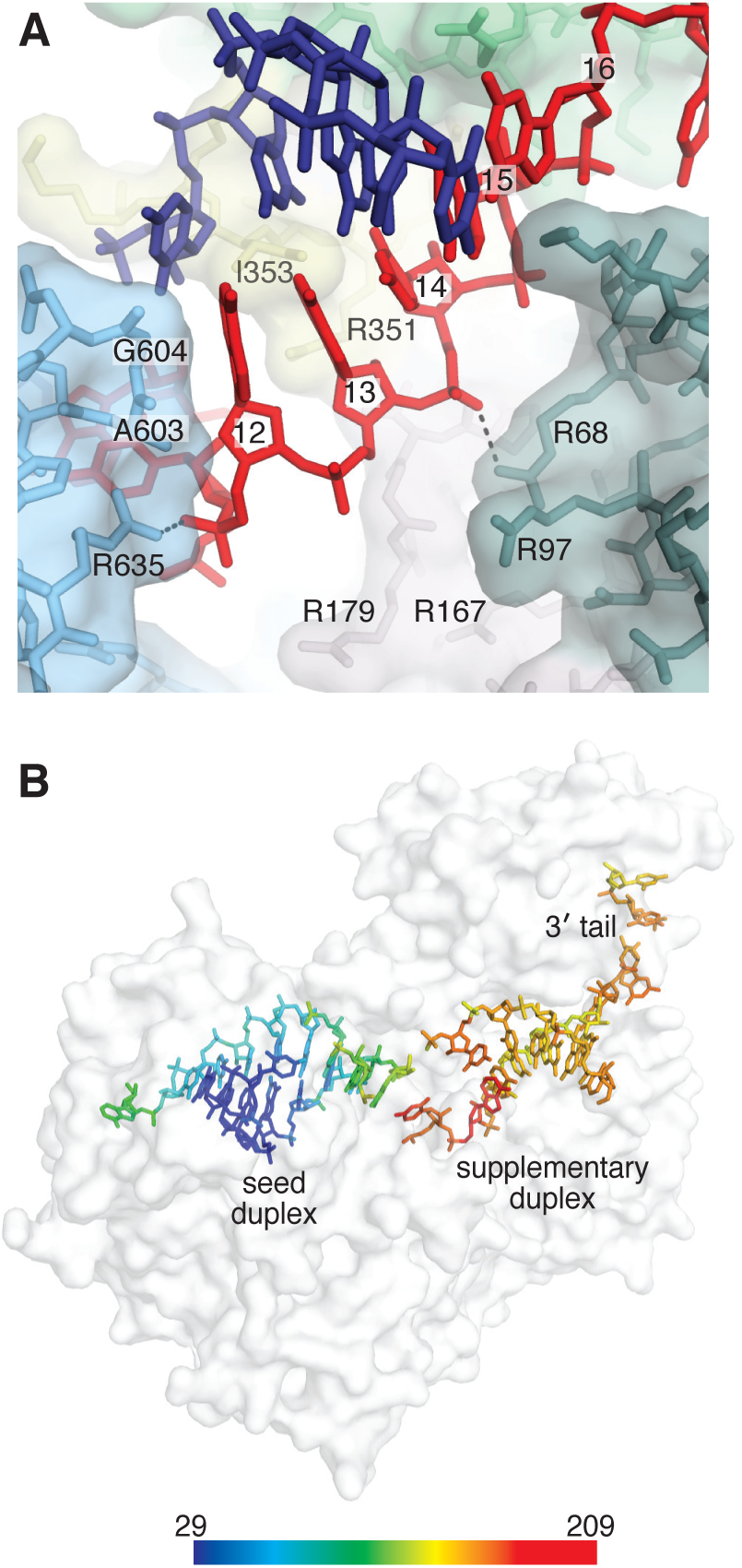
The supplementary duplex is mobile. **A.** Direct contacts to the supplementary miRNA-target duplex shown. Dashed lines indicate potential salt-linkages. **B.** Protein shown as semi-transparent surface representation with miRNA and target colored by temperature factor (scale shown at bottom).

## Discussion

Seminal structural studies of *T. thermophilus* Argonaute led to a model in which guide-target pairing initiates in the seed region and then propagates, in a direct fashion, towards the guide 3′ end (Wang, Juranek et al., 2009). However, the applicability of this model in mammalian Argonaute systems has recently been questioned (Bartel, 2018). Our work here provides evidence that human Ago2 may use a distinct mechanism for target recognition beyond the seed. The observation that Ago2 crystallized in a conformation that avoided central pairing, despite the presence of complementarity to that region, not only demonstrates that Ago2 can make supplementary interactions without central pairing, but also suggests that this may be an energetically favored route for establishing miRNA-target pairing beyond the seed.

Based on our results, we suggest that supplementary interactions may serve as an initial nucleation site for 3′ and/or central pairing. In this model, Ago2 would use the supplementary region to interrogate seed-bound target RNAs for potential 3′ end complementarity prior to committing to the costly step associated with opening the central cleft, loss of 3′ end association, and guide:target winding. To accommodate further pairing within the central region, the central cleft would have to open more substantially than observed here. Notably, this would coincide with increased tension in the miRNA 3′ tail, as it would induce movement of the PAZ domain away from the central cleft. This interplay suggests a critical mechanism employed by Argonaute in the regulation of duplex progression: conformational shift of the protein may be coupled to tension in the miRNA, arising from the anchoring of both the 5′ and 3′ ends within the MID and PAZ domains, respectively. Therefore, pairing in the central region would require not only opening of the protein, but also loss of 3′ end retention. A similar model for miRNA target recognition and pairing propagation was recently proposed (Bartel, 2018). In support of the model, it has been shown that endonucleolytic cleavage of a centrally-paired target RNA requires complementarity extending to positions g16 or g17 (Ameres et al., 2007, De et al., 2013, Haley & Zamore, 2004). The central nucleotides g9-g12, moreover, contribute minimally to guide:target affinity and association (Grosswendt et al., 2014, Salomon et al., 2015, Wee et al., 2012). And finally, target-binding and single molecule assays reveal a reproducible reduction in target affinity upon pairing to g9 across multiple miRNA sequences (Salomon et al., 2015, Schirle et al., 2014).

Taken together, these findings indicate that opening of the central cleft, coupled with the requisite release of the miRNA 3′ end, presents a substantial energetic barrier associated with central pairing and guide:target winding. Here, we suggest that central pairing is licensed by opening of the PIWI Loop and L2 Loop, forming a “central gate” within the central RNA binding cleft. This mechanism may explain why the mammalian Argonautes have evolved away from slicing as the predominant form of target silencing and, moreover, would ensure that the protein does not employ its endonucleolytic activity spuriously. Notably, the crystalized target RNA used here did not contain pairing to the miRNA 3′ tail region (g17-g21), which is important for specific miRNA-target interactions in vivo (Broughton, Lovci et al., 2016a) and the process of target directed miRNA degradation (TDMD) (Fuchs Wightman, Giono et al., 2018). Biochemical data also indicate Ago2 is generally capable of using the 3′ tail for target recognition (McGeary, Lin et al., 2018). Considering the size of the supplementary chamber, which can only accommodate 5 miRNA:target base pairs, we suggest that the Ago2-miRNA complex must adopt a unique conformation, distinct from the seed plus supplementary paired structure, to engage targets with extended 3′ pairing.

Our findings also provide mechanistic insights for improved understanding of supplementary miRNA-target interactions. The finding that a GC-rich supplementary sequence can increase target affinity >20-fold, reveals that supplementary interactions can influence target recognition to a greater extent than is widely appreciated (Agarwal et al., 2015, Bartel, 2018, Wee et al., 2012). Moreover, the observations that, under specific circumstances: 1) seed and supplementary regions can be effectively bridged by gaps 15 nt in length; and, 2) the presence of even just two GC supplementary base pairs can significantly increase target affinity; raise the possibility that some supplementary sites may not be easily recognized, and thus may often go undetected. Indeed, analysis of 477 sequences in miRBase (Kozomara & Griffiths-Jones, 2011) reveals that about 25% of human miRNAs contain ≥3 GC nucleotides within positions g13–g16.

Finally, the finding that increases in miRNA 3′ tail length substantially increase the stability of supplementary interactions and potentially expand the miRNA nucleotides available for target recognition indicate that isomiR form may be an important consideration when predicting which nucleotides in the 3′ tail are able to make supplementary interactions as well as the overall contribution of supplementary pairing to target recognition. To our knowledge, these results also provide the first direct evidence that target recognition by the Ago2-miRNA complex can be tuned by 3′ isomiR length. Considering the regulation and abundance of diverse isomiR forms in animals (Fernandez-Valverde, Taft et al., 2010, Wyman, Knouf et al., 2011, Yu, Pillman et al., 2017), we suggest that control of 3′ isomiR length may be a widespread mechanism for modulating supplementary interactions and tuning miRNA target specificity.

## Materials and Methods

### Oligonucleotides

<u>miR-122 variants</u>

miR-122-21mer: 5′-Phosphate-rUrGrGrArGrUrGrUrGrArCrArArUrGrGrUrGrUrUrU-3′

miR-122-22mer: 5′-Phosphate-rUrGrGrArGrUrGrUrGrArCrArArUrGrGrUrGrUrUrUrG-3′

miR-122-23mer: 5′-Phosphate

rUrGrGrArGrUrGrUrGrArCrArArUrGrGrUrGrUrUrUrGrU-3′

miR-122-strong-sup: 5′-Phosphate

rUrGrGrArGrUrGrUrGrArCrArGrCrGrGrCrGrArArArG-3′

miR-122-weaker-sup: 5′-Phosphate

rUrGrGrArGrUrGrUrGrArCrArCrUrArUrCrGrArArArG-3′

<u>miR-122-strong-sup-passenger: 5’</u>

<u>rUrUrCrGrCrCrGrCrUrArUrCrArCrArCrUrArArArUrA-3’</u>

<u>miR-122 purification oligonucleotides</u>

Capture oligo: 5′-Biotin-mUmCmUmCmGmUmCmUmAmAmCmCmAmUmGmCmCmAmAmCmAmC mUmCmCmAmAmCmUmCmU-3′

Competitor DNA: 5′-Biotin-GCAGAGATCAAGTGTTCGCATGGTTAGCAGAGA-3′

<u>miR-122 target RNAs</u>

2–16 perfect: 5′-rCrCrArUrUrGrUrCrArCrArCrUrCrCrArArA-3′

seed-only: 5′-rArArArArArArArArArCrArCrUrCrCuAA-3′

strong sup, 1nt bridge: 5′-rCrCrGrCrArArCrArCrUrCrCrUrArA

strong sup, 2nt bridge: 5′-rCrCrGrCrArArArCrArCrUrCrCrUrArA-3′

strong sup, 4nt bridge: 5′-rCrCrGrCrArArArArArCrArCrUrCrCrUrArA-3′

strong sup, 10nt bridge: 5′-"CrCrGrCrArArArArArArArArArArArCrArCrUrCrCrUrArA-3′

strong sup, 15nt bridge: 5′-"CrCrGrCrArArArArArArArArArArArArArArArArCrArCrUrCrCrUrArA-3′

weak sup, 1 nt bridge: 5′-"ArUrArGrArArCrArCrUrCrCrUrArA-3′

weak sup, 2 nt bridge: 5′-"ArUrArGrArArArCrArCrUrCrCrUrArA-3′

weak sup, 4 nt bridge: 5′-"ArUrArGrArArArArArCrArCrUrCrCrUrArA-3′

weak sup, 10 nt bridge: 5′-"ArUrArGrArArArArArArArArArArArCrArCrUrCrCrUrArA-3′

2–8 plus 13–17: 5′-"ArCrGrCrArArCrCrArUrGrCrCrArArCrArCrUrCrCrArArA-3′

<u>Luciferase reporter assay target sites cloning primers</u>

seed-only-f: 5’-TCGAGAAAAAAAAAAAAAAACACTCCTAAGC-3’

seed-only-r: 5’-GGCCGCTTAGGAGTGTTTTTTTTTTTTTTTC-3’

1nt-bridge-f: 5’-TCGAGAAAAAAAAACCGCAACACTCCTAAGC-3’

1nt-bridge-r: 5’-GGCCGCTTAGGAGTGTTGCGGTTTTTTTTTC-3’

2nt-bridge-f: 5’-TCGAGAAAAAAAACCGCAAACACTCCTAAGC-3’

2nt-bridge-r: 5’-GGCCGCTTAGGAGTGTTTGCGGTTTTTTTTC-3’

4nt-bridge-f: 5’-TCGAGAAAAAACCGCAAAAACACTCCTAAGC-3’

4nt-bridge-r: 5’-GGCCGCTTAGGAGTGTTTTTGCGGTTTTTTC-3’

10nt-bridge-f: 5’-TCGAGCCGCAAAAAAAAAAACACTCCTAAGC-3’

10nt-bridge-r: 5’-GGCCGCTTAGGAGTGTTTTTTTTTTTGCGGC-3’

15nt-bridge-f: 5’-TCGAGCCGCAAAAAAAAAAAAAAAACACTCCTAAGC-3’

15nt-bridge-r: 5’-GGCCGCTTAGGAGTGTTTTTTTTTTTTTTTTGCGGC-3’

20nt-bridge-f: 5’-TCGAGCCGCAAAAAAAAAAAAAAAAAAAAACACTCCTAAGC-3’

20nt-bridge-r: 5’-GGCCGCTTAGGAGTGTTTTTTTTTTTTTTTTTTTTTGCGGC-3’

### Preparation of Ago2-miRNA-122 complexes

Human Ago2 loaded with miR-122 variants was purified as described previously (Schirle et al., 2014). Briefly, His_6_-tagged Ago2 was expressed in Sf9 cells using a baculovirus system (Invitrogen). Cells were harvested by centrifugation and lysed in Lysis Buffer (300mM NaCl, 0.5 mM TCEP, 50 mM Tris, pH 8) using a single pass through a M-110P lab homogenizer (Microfluidics). Lysate was cleared and the soluble fraction was applied to Ni-NTA resin (Qiagen) and incubated at 4°C for 1.5 h in 50 ml conical tubes. Resin was pelleted by brief centrifugation and the supernatant solution was discarded. The resin was washed with ~50 ml ice cold Nickel Wash Buffer (300 mM NaCl, 300 mM imidazole, 0.5 mM TCEP, 50 mM Tris, pH 8). Centrifugation/wash steps were repeated a total of three times. Co-purifying cellular RNAs were degraded by incubating with micrococcal nuclease (Clontech) on-resin in ~15 ml of Nickel Wash Buffer supplemented with 5mM CaCl_2_ at room temperature for 45 minutes. The resin was washed three times again with Nickel Wash Buffer and then eluted in four column volumes of Nickel Elution Buffer (300 mM NaCl, 300 mM imidazole, 0.5 mM TCEP, 50 mM Tris, pH 8). Eluted Ago2 was incubated with a synthetic miR-122 variant and TEV protease (to remove the N-terminal His_6_ tag on recombinant Ago2) during an overnight dialysis against 1–2 liters of Hi-Trap Dialysis Buffer (300 mM NaCl, 15 mM imidazole, 0.5 mM TCEP, 50 mM Tris, pH 8) at 4°C. The dialyzed protein was then passed through a 5 ml Hi-Trap Chelating column (GE Healthcare Life Sciences) and the unbound material was collected. The Ago2 molecules loaded with miR-122 were purified using a modified Arpon method (Flores-Jasso, Salomon et al., 2013). Loaded molecules were further purified by size exclusion using a Superdex 200 Increase 10/300 column (GE Life Sciences) in High Salt SEC Buffer (1M NaCl, 0.5mM TCEP, 50mM Tris pH 8). The final protein was dialized into Tris pH 8 Crystal Buffer (10mM Tris pH 8, 100mM NaCl), concentrated to ~8 mg/ml, aliquoted and flash frozen and stored at −80°C. Samples were thawed slowly on ice for all experiments.

### Crystallization and diffraction data collection

Crystallization samples were prepared by diluting purified Ago2-miR-122 to a final concentration of 1 mg/mL and adding 2–16 perfect target RNA in a 1:1.2 molar ratio. Adding target directly to concentrated Ago2-miR-122 would often result in precipitation and was therefore avoided. The sample was then incubated briefly on ice before used for screening. Crystals were grown by hanging drop vapor diffusion at 20°C. Initial hits were obtained in the JBScreen Classic screen (Jena Bioscience), which included a condition that grew crystals as clusters of needles. These needles were grown in drops containin 16% PEG 6000 and 10mM tris-Sodium citrate (condition 4/B2) and were reticent to optimization with similar chemical regimes. Needles were therefore harvested and crushed into micro-seeds using a Seed Beed (Hampton Research) to be used in a wider screening effort. New conditions were screened by mixing seeds, ternary Ago2 complex, and new conditions in a 0.2:1:1 ratio in hanging drops. Larger, more rod-like crystals were obtained in a new condition, and iterative screening and optimization in this condition (26% MPD, 5% PEG 3350, 50 mM imidazole pH 8) resulted in crystals that were well-ordered in the center but grew frayed at the edges. Crystals were harvested directly in nylon loops and flash frozen in liquid nitrogen. Diffraction data were collected under cryogenic conditions remotely at beam lines 12-2 at the Stanford Synchrotron Radiation Lightsource (SSRL) using a micro-focus beam to avoid the multiple lattices at the edges. Data were processed using XDS and Scala (Kabsch, 2010, Winn, Ballard et al., 2011).

### Model building and refinement

Ago2-miR122-target ternary complex structure was solved by molecular replacement using the MID-PIWI lobe and the PAZ and N-domains of the seed-paired Ago2 structure (4W5O) as sequential search models with Phaser-MR in the PHENIX graphical interface (McCoy, Grosse-Kunstleve et al., 2007). Electron density for RNA in the seed and supplementary chambers was observed in the molecular replacement map. Atomic models were built using Coot and submitted to XYZ coordinate, TLS, and B-factor refinement using PHENIX (Adams, Afonine et al., 2010). Model building and refinement were carried out iteratively until all interpretable electron density was modeled. All structure figures were generated using PyMOL (Schrödinger, LLC) or Chimera (UCSF).

### Equilibrium target binding assays

Equilibrium dissociation constants were determined as described previously (Schirle et al., 2014). Briefly, various concentrations of the Ago2-miR-122 complex were incubated with 0.1 nM ^32^P 5′-radiolabeled target RNA in binding reaction buffer (30 mM tris pH 8.0, 100 mM potassium acetate, 2 mM magnesium acetate, 0.5 mM TCEP, 0.005% (v/v) NP-40, 0.01 mg/mL baker’s yeast tRNA), in a reaction volume of 100 µL at room temperature for 60 min. Using a dot-blot apparatus (GE Healthcare Life Sciences), protein-RNA complexes were captured on Protran nitrocellulose membrane (0.45 µm pore size, Whatman, GE Healthcare Life Sciences) and unbound RNA on Hybond Nylon membrane (Amersham, GE Healthcare Life Sciences). Samples were applied with vacuum and then washed one with 100 µL of ice-cold wash buffer (30 mM Tris pH 8.0, 0.1 M potassium acetate, 2 mM magnesium acetate, 0.5 mM TCEP). Membranes were air-dried and ^32^P signal was visualized by phosphorimaging. ImageQuant was used to quantify data and dissociation constants calculated using Prism version 6.0g (GraphPad Software, Inc.), with the following formula, which accounts for potential ligand depletion (Wee et al., 2012):

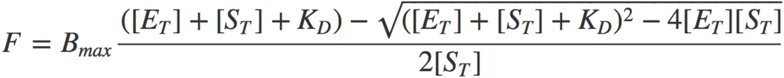

where *F* = fraction of target bound, *B*_*max*_ = calculated maximum number of binding sites, *[E*_*T*_] = total enzyme concentration, *[S*_*T*_] = total target concentration, and *K*_*d*_ = apparent equilibrium dissociation constant.

### Target release assays

Target release assays were performed by incubating purified Ago2-miR-122 complexes at a final concentration of 2 nM with 0.1 nM of ^32^P 5′-radiolabeled target RNA in binding reaction buffer (30 mM tris pH 8.0, 100 mM potassium acetate, 2 mM magnesium acetate, 0.5 mM TCEP, 0.005% (v/v) NP-40, 0.01 mg/mL baker’s yeast tRNA) in a final volume of 700 µl. Samples were allowed to come to equilibrium by incubating at room temperature for 45 minutes. The time point 0 aliquot (50 µl) was removed, and bound and free RNA were separated using a dot-blot apparatus (GE Healthcare Life Sciences). A small volume (6.5 µl) of concentrated unlabeled target RNA was then added to a final concentration of 100 nM. Further samples were removed at various time points and separated as before. ^32^P-labeled target RNA was visualized and quantified by phosphorimaging, as described above. The fraction of the ^32^P-labeled target bound to Ago2 was plotted as a function of time, normalized to timepoint 0, and fit to a first order exponential using Prism version 6.0g (GraphPad Software, Inc.).

### Luciferase reporter assays

Target sites were cloned into psiCHECK2 plasmid (Promega Corporation) using Xho1 and Not1 sites. The miRNA duplex was made by denaturing 1:1 mixture of guide and passenger strands in annealing buffer (10mM Tris pH8.0, 50mM NaCl) at 95°C for 2 minutes, then cooling to room temperature by placing on the benchtop. Luciferase reporter assays were performed with HEK 293 cells. The HEK 293 cells were plate at 10^4^ cells with 80ul DMEM complete media per well in 96-well plate (Thermal Scientific, Cat#136101), and cultured in a temperature, CO2 and humidity controlled cell incubator for ~4h to allow cells attach to the bottom of the plate. Cells in each well were transfected with 40 ng psiCHECK2 plasmid and titration of miRNA duplex (0, 1.56, 3.12, 6.25, 12.5, 25 nM) using lipofectamine 2000 (Invitrogen) based on manufacturer’s protocol. Each transfection condition was prepared at least in triplicate. After a 24 hour incubation, media in each well was removed, and luciferase activities were measured using Dual-Glo^®^ Luciferase Assay System (Promega Corporation, Cat#E2920) based on manufacturer’s description. The ratio of Renilla signal to Firefly signal was normalized to 0 nM miRNA transfection condition, and analyzed using Prism version 6.0g (GraphPad Software, Inc.), with the one-phase decay formula:

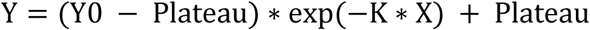

Where Y is normalized Renilla/Firefly, X is the concentration of miRNA duplex transfected, Y0 is the Renilla/Firefly when miRNA is 0 nM, Plateau is the limit of the Renilla/Firefly ratio as [miRNA] approaches infinity. Fold repression is reported as the plateau value of a no target site control divided by the plateau value of a construct bearing a target site.

## Author Contributions

J.S.G. purified Ago2-miRNA samples, conducted crystallization experiments, x-ray diffraction data collection, structural determination and refinement, and structural interpretation, and wrote the manuscript. Y.X. purified Ago2-miRNA samples, and designed and performed equilibrium binding experiments, and luciferase reporter experiments. L.F.R.G. purified Ago2-miRNA samples, and designed and performed 3′ isomiR off-rate experiments. I.J.M. advised experiments and revised the manuscript.

## Acknowledgements

We are grateful to S. M. Klum and T. Anzelon for support and thoughtful discussions J.S.G. was a Pre-doctoral Fellow of the American Heart Association and an Abrams Charitable Trust Award recipient. L.F.R.G. is supported by the Swiss National Science Foundation [Advanced Postdoc Mobility Fellowship P300PA_177860]. I.J.M. and this research were supported by NIH grants GM104475, GM115649, and GM127090. Diffraction data were collected at beam line 12-2 at the Stanford Synchrotron Radiation Lightsource. Coordinates of the seed plus supplementary-paired Ago2-miRNA-target complex have been deposited in the Protein Data Bank (PDB) (6N4O).

---

**Fig. EV1.**
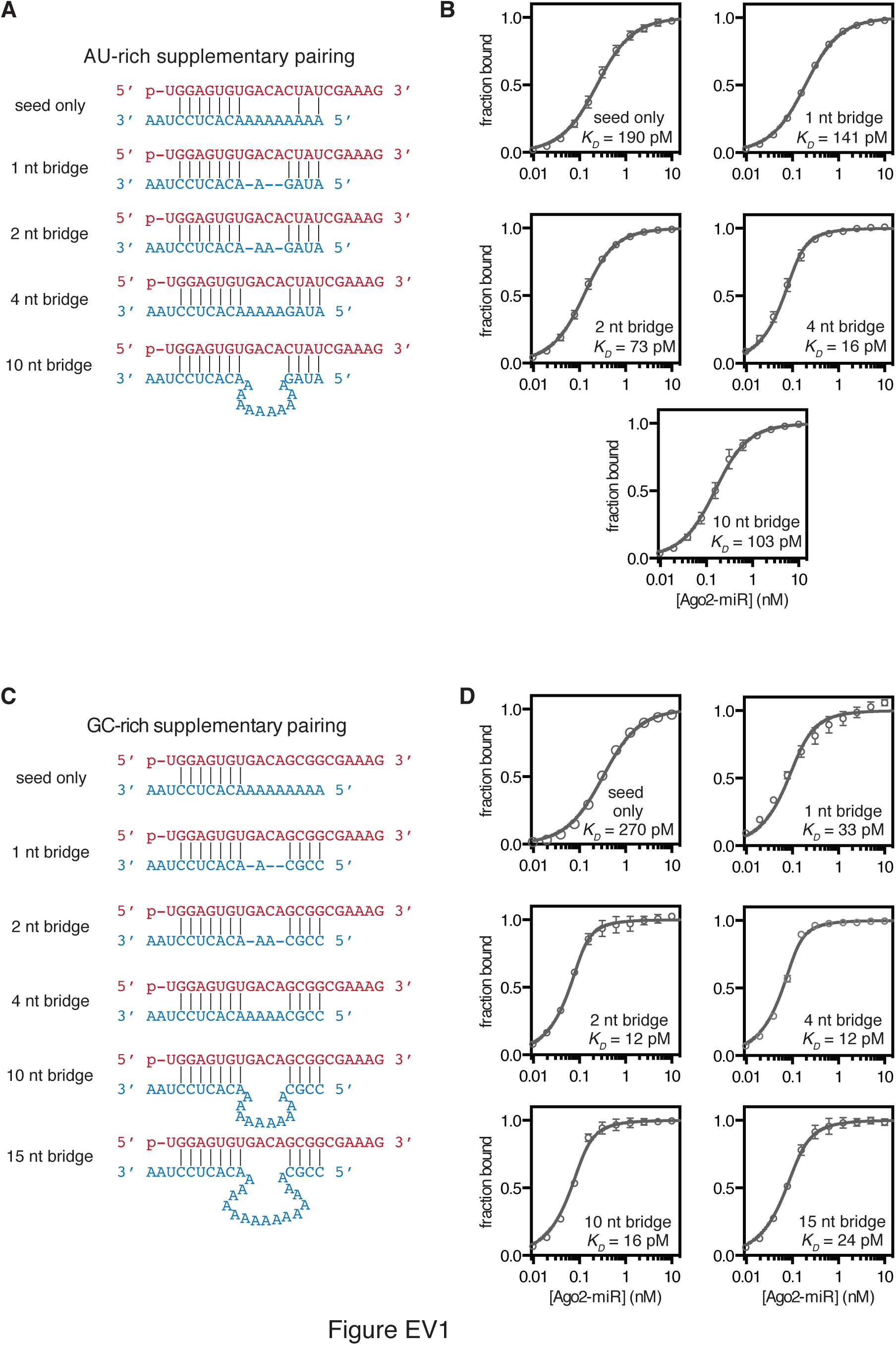
Binding data for analysis of seed-supplementary bridging region. **A.** Variant-1 of miRNA-122 (with an AU-rich supplementary region) shown paired to target RNAs containing bridging regions of various lengths. **B.** Fraction target bound at equilibrium versus Ago-miRNA concentration. Calculated *K*_*d*_ values indicated. **C.** Variant-2 of miRNA-122 (with GC-rich supplementary region) shown paired to target RNAs containing bridging regions of various lengths. **D.** Fraction target bound at equilibrium versus Ago-miRNA concentration. Calculated *K*_*d*_ values indicated. All plotted data are the means of at least three independent replicates. Error bars indicate SEM.

**Fig. EV2.**
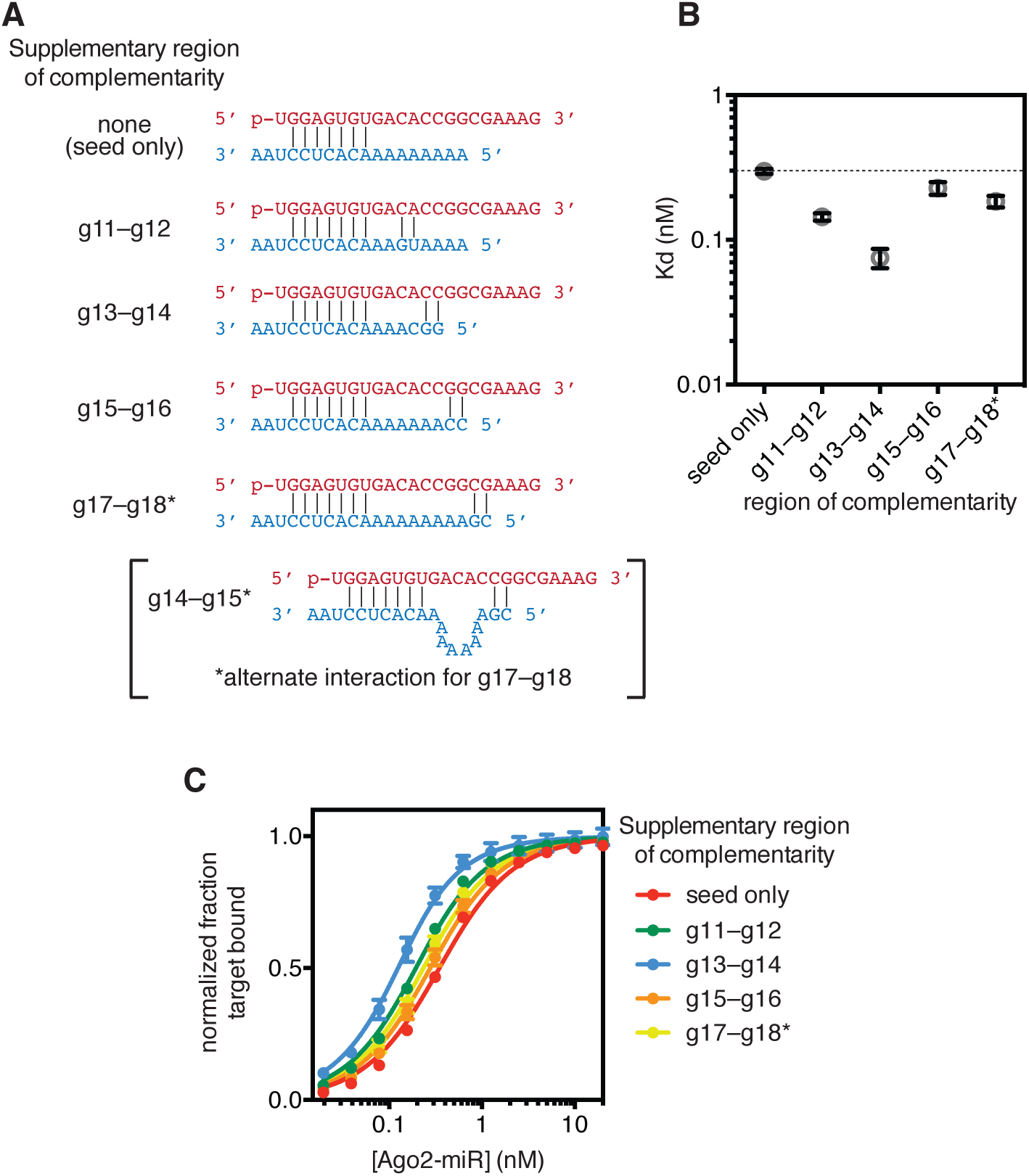
Analysis of di-nucleotide supplementary pairing. **A.** Variant-2 of miRNA-122 (red) shown paired to target RNAs (blue) with supplementary interactions restricted to two adjacent base pairs. Two modes of interaction are possible for the g17-g18 target. **B.** Dissociation constants (*K*_*d*_) of the Ago2-miRNA complex binding to target RNAs in panel A. C. Binding curves used to calculate *K*_*d*_ values in panel B. All plotted data are the means of at least three independent replicates. Error bars indicate SEM.

**Fig. EV3.**
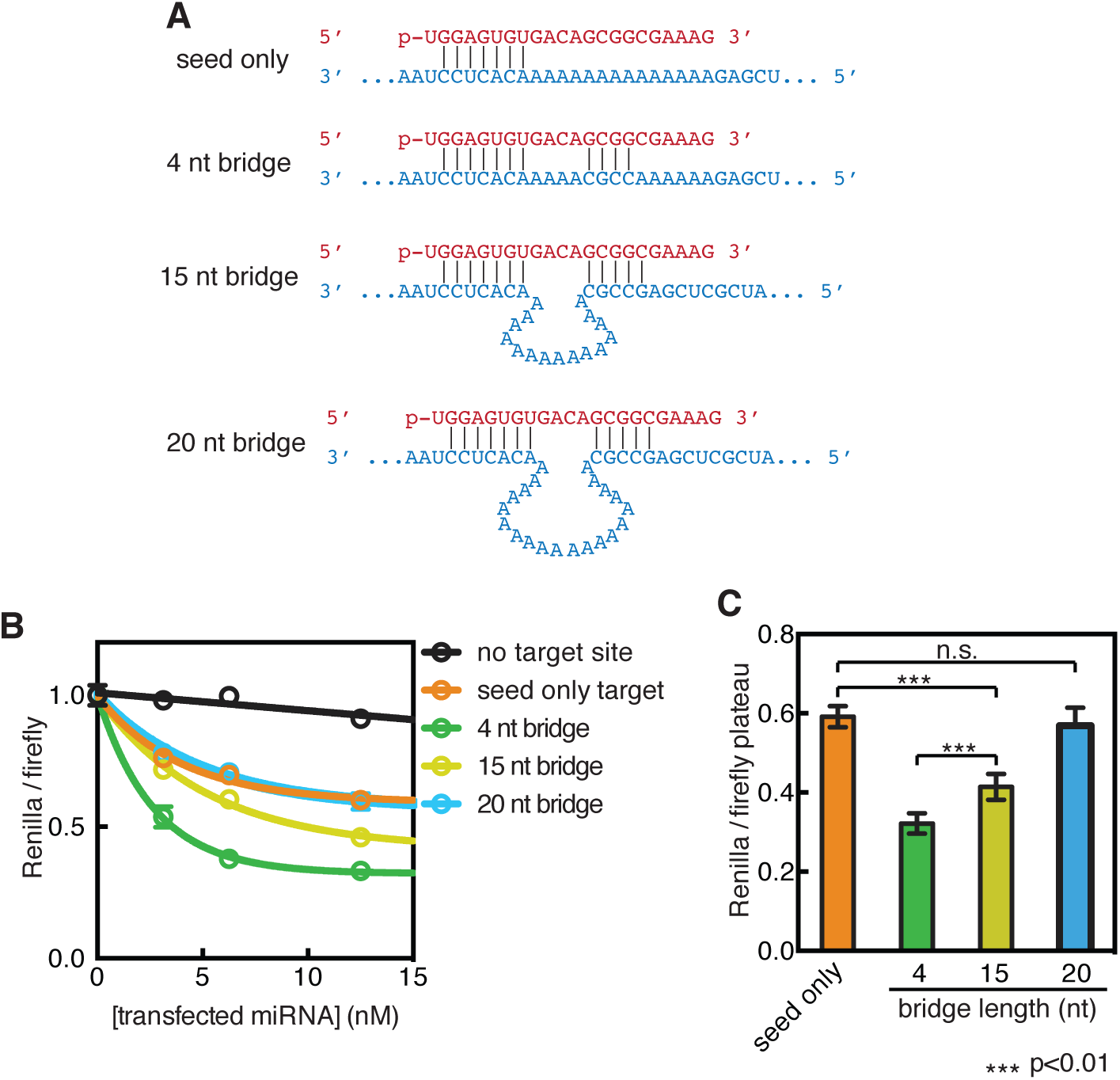
Impact of bridging region on target-reporter repression. **A.** Variant-2 of miRNA-122 (red) shown paired to single target sites (blue) cloned into the 3′ UTR of a Renilla luciferase reporter. **B.** Renilla luciferase levels (normalized to Firefly internal control) plotted as a function of miRNA concentration co-transfected into HEK 293 cells. **C.** Plateau values from panel B for targets with bridging regions of various lengths. All plotted data are the means of at least three biological replicates. Error bars indicate SEM.

